# Effects of periodic group housing opportunities on reproductive performance and welfare in sows

**DOI:** 10.64898/2026.05.19.726187

**Authors:** Tomoya Shimasaki, Ken-ichi Yayou, Tomoki Kojima, Chen-Yu Huang, Hiromi Kato, Mitsuyoshi Ishida, Ken-ichi Takeda

## Abstract

**Objective:** Stall housing of pregnant sows raises welfare concerns, whereas conventional group housing systems often reduce space efficiency. This study evaluated the effects of periodic group housing (PG) on reproductive performance and welfare compared with continuous stall housing (CS).

**Methods:** Sows in the CS group (n = 15) were continuously housed in stalls. In the PG group (n = 15), sows were housed in groups of three and allocated 1 day of group housing and 6 days of stall housing per week over 10 weeks. During group housing sessions, the sows had access to a group housing area containing sawdust. Behavioral observations and salivary cortisol measurements were conducted on the first day of the stall housing session in weeks 1, 6, and 10. Behavioral indices were expressed as proportions based on 90 sampling points recorded at 1-min intervals.

**Results:** The number of stillbirths was significantly lower in the PG group than in the CS group (0.63 vs. 1.49 piglets per litter). whereas other reproductive outcomes, including total litter size and average birth weight, did not differ. In older parity sows, the PG treatment markedly increased the proportion of time spent lying, suggesting reduced discomfort associated with restricted movement. Furthermore, the proportion of exploratory behavior decreased markedly, and drinking behavior showed a decreasing trend across parity levels in the PG group, suggesting partial satisfaction of motivations for environmental exploration and oral manipulation. The proportion of oral abnormal behavior showed a pronounced interaction between housing treatment and experimental week, increasing from week 1 to week 6 in the PG group. Salivary cortisol concentrations did not differ between the groups.

**Conclusion:** PG may improve reproductive performance and partially satisfy the behavioral motivations restricted under continuous stall housing. This system may represent a practical alternative for improving animal welfare while minimizing economic losses.

## INTRODUCTION

In the modern swine industry, stall housing for pregnant gilts and sows has raised animal welfare (AW) concerns due to its severe restriction of movement. Consequently, a transition toward group housing systems has progressed, particularly within the European Union (EU). Group housing systems enable social interaction and exploratory behavior, both of which are innate behaviors in domesticated pigs. In contrast, pigs housed in stalls are unable to express these behaviors, which have been associated with adverse effects on behavioral stress indicators, including increased standing time, reduced lying time [1,2], and a higher frequency of oral abnormal behaviors, such as sham-chewing and bar-biting [1,3,4].

However, conventional group housing systems present several economic challenges. First, the capital investment required to convert from stall housing to group housing systems is substantial. This concern was among the primary barriers to the adoption of group housing in Belgium before the implementation of EU legislation banning stall housing [5]. Additionally, group housing systems require more space and often result in fewer animals per unit area [6,7], which remains a key factor delaying the widespread adoption of improved AW systems. Indeed, according to a questionnaire survey of pig farmers in Japan, typical concerns for “management in consideration of AW” are increases in production costs and decreases in farm productivity [8]. Therefore, there is a need for practical alternative housing systems that improve both AW and farm economic viability, particularly in Asian regions, where swine production is most prevalent globally. One potential management strategy is to provide pregnant sows with periodic opportunities for individual exercise outside the stall. Previous studies have reported that periodic exercise, involving walking along corridors within the swine facility, has positive physiological and reproductive effects, including improved umbilical cord blood flow [9], increased bone density and the number of weaned piglets [10], and a higher number of live-born piglets in older sows [11]. However, this form of individual exercise management is not practically feasible, as it requires long-term assistance from stockpersons. Furthermore, it does not provide opportunities for social contact.

Therefore, we propose a management system in which stall-housed pregnant sows are given access to a group housing area in small groups at periodic intervals. This system allows pregnant sows to exercise, explore outside the stalls, and engage in limited social interaction while requiring less assistance from stockpersons. Additionally, compared with conventional group housing systems, the proposed system requires less space and may be a more feasible option for farmers concerned about economics. However, to our knowledge, no studies have evaluated the effects of such a “Periodic group (PG) housing system” on the reproductive performance, behavioral outcomes, and physiological stress in pigs. Hence, in this study, we evaluated the effects of the proposed system on reproductive performance, behavior, and physiological stress indicators.

## MATERIALS AND METHODS

All experimental procedures were conducted in accordance with the Guide for the Care and Use of Experimental Animals and were approved by the Animal Care Committee of the National Institute of Livestock and Grassland Science (Approval No.: R5-P05-NILGS). The experiments were conducted between August 15, 2023, and July 18, 2024.

### Animals and housing treatment

Twenty-six pregnant sows of a hyper-prolific Landrace × Large White crossbreed (ZEN-NOH HI-COOP; ZEN-NOH Livestock Co., Ltd., Tokyo, Japan) were used in this study. Four sows were examined over two parities; therefore, data were collected from 30 gestational periods (parity range: 1–5; mean parity: 2.2 ± 1.2). The animals were fed once daily at 09:00 h, as recommended by the breeding company.

Before the experiment, all sows had been kept in either individual gestation stalls or farrowing crates and had not experienced group housing for at least 10 weeks. After pregnancy confirmation on days 24–27 post-artificial insemination, sows were assigned to one of two treatments: continuous stall housing (**CS**, n = 15) or periodic group housing (**PG**, n = 15). The assignment was balanced with respect to parity and season. In the PG treatment, sows were managed in groups of three, resulting in five groups in total. Sows in the CS treatment were maintained in standard individual stalls (0.7 × 1.8 m^2^) throughout gestation. Sows in the PG group were provided with 1 day of group housing per week and 6 days of stall housing over 10 weeks (Fig. 1). During group housing sessions, sows had access to both their home stalls and an adjacent group housing area (total area: 13.8 m^2^). Each stall was equipped with a nipple drinker, and approximately 5 mm of sawdust bedding was provided in the group housing area. Group housing sessions commenced after all sows in a group had completed feeding and ended before feeding the following morning. All sows were moved to individual farrowing crates 1 week before the expected farrowing date.

**Figure 1.**
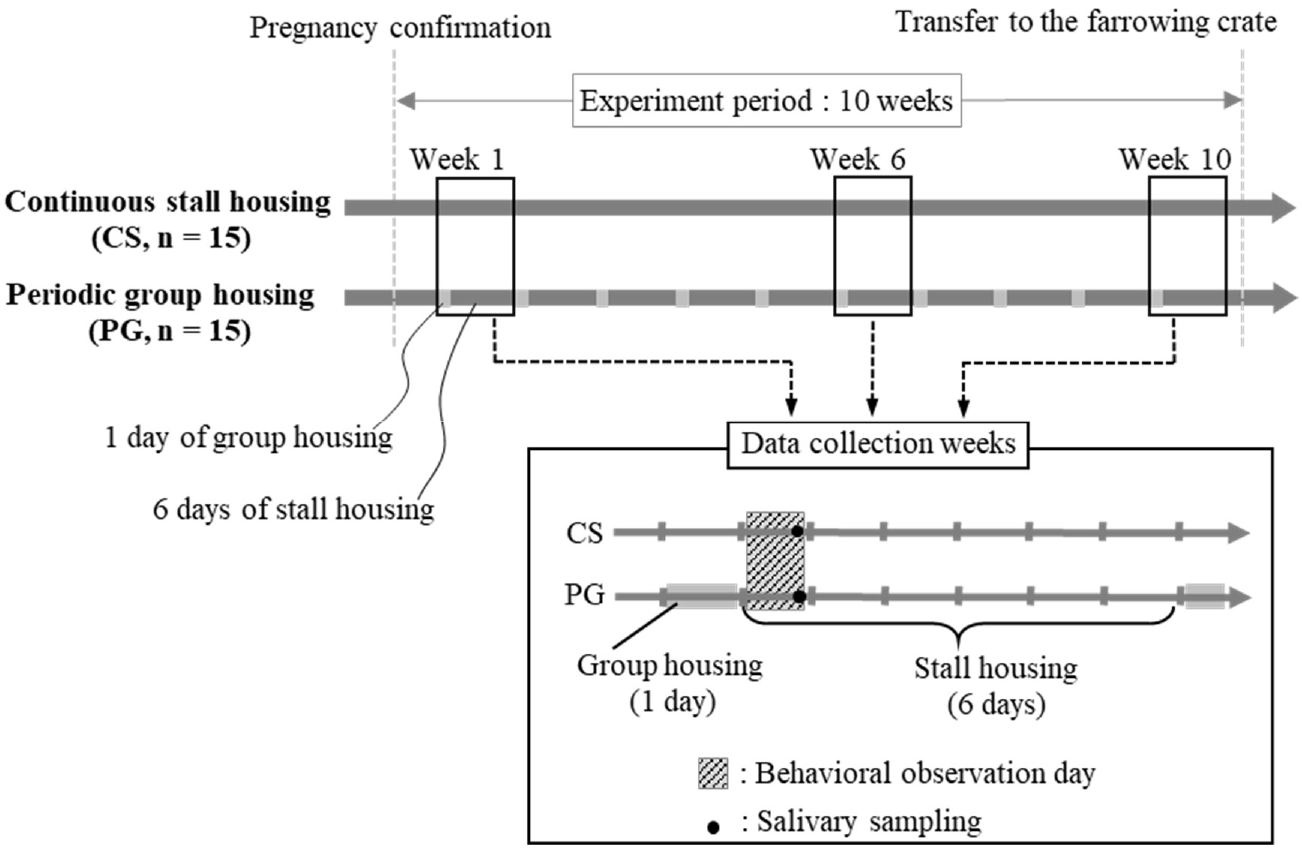
Experimental timeline of sows assigned to continuous stall housing (CS, n = 15) or periodic group housing (PG, n = 15) during the 10-week period after pregnancy confirmation. Sows in the PG treatment underwent repeated cycles consisting of 1 day of group housing followed by 6 days of stall housing. Behavioral observations were conducted on the second day of weeks 1, 6, and 10 (shaded area). Saliva samples for cortisol analysis were collected on the following mornings (black dots). All sows were moved to individual farrowing crates 1 week before the expected farrowing date.

### Data collection

To evaluate reproductive performance, total litter size was recorded. This included the sum of stillbirths, neonatal mortality (defined as deaths occurring before the onset of suckling due to crushing or weakness), and live piglets at the beginning of lactation. Average birth weight was also measured.

Throughout gestation, behavioral data were continuously recorded during the experimental period using surveillance cameras (ZOSI Technology Ltd., Zhuhai, China) positioned 2.1 m in front of the pens at a height of 2.0 m. Behavioral observations were conducted using recordings from the first day of the stall housing sessions in weeks 1, 6, and 10. Each observation day lasted 90 min, consisting of six 15-min observation periods at 11:00, 12:00, 13:00, 15:00, 16:00, and 17:00. These time points were selected to avoid the potential effects of feeding and cleaning activities on behavior. Definitions of each behavioral category are provided in the ethogram (Table 1). Postural behaviors (standing, lying, and sitting) were recorded using scan sampling, whereas behavioral events (drinking, exploration, and oral abnormal behavior) were recorded using one-zero sampling. Both sampling methods were conducted at 1-min intervals. Behavioral data were summed for each observation day for statistical analysis.

**Table 1.** Ethogram for behavioral observation.

| Behavioral index | Definition | Recording method |
| --- | --- | --- |
| Standing | Standing on all four legs | Scan sampling |
| Lying | Lying in sternal or lateral recumbency |  |
| Sitting | Sitting posture with forelimbs extended and body weight supported on the rump. |  |
| Drinking | Drinking water, pushing the nipple drinker, or holding it in the mouth | One-zero sampling |
| Exploring | Rooting or licking the floor surface or trough with head shaking |  |
| Oral abnormal behavior | Chewing with no object in the mouth or nosing, rubbing, or biting any metal bar of the stall |  |

Saliva samples for cortisol analysis were collected between 08:30 and 09:00 h on the second day of the stall housing session in weeks 1, 6, and 10. Sows were allowed to chew on absorbent cotton held with forceps. Saliva was then extracted from the cotton by centrifugation in a collection tube (Swab Storage Tube; Salimetrics, LLC., California, United States) at 2,000 × *g* for 20 min. The supernatant saliva was transferred into 1.5 mL plastic tubes and stored at −30°C until analysis. Cortisol concentrations were measured in duplicate using a cortisol ELISA kit (ADI-901-071; Enzo Biochem, Inc., New York, United States) according to the manufacturer’s instructions. Optical density was measured at 405 nm using a microplate reader (Infinite50; TECAN Group Ltd., Männedorf, Switzerland).

### Statistical analysis

All statistical analyses were conducted using R (version 4.5.3) [12]. Statistical significance was set at *p* = 0.05, and *p*-values between 0.05 and 0.10 were considered indicative of a statistical trend.

For analysis, sows were classified into three parity groups based on individual parity records: Young (parity 1), Middle (parity 2–3), and Old (parity 4–5). The distribution of sows across treatments and parity groups was as follows: CS—Young: n = 5, Middle: n = 7, Old: n = 3; PG—Young: n = 7, Middle: n = 6, Old: n = 2.

Reproductive performance data, except for average birth weight, were analyzed using generalized linear mixed models (GLMMs) with a Poisson distribution and a log link function. Average birth weight was analyzed using a linear mixed model (LMM). All models included housing treatment and parity group as fixed effects and animal as a random effect. Initial models also included the interaction between housing treatment and parity group; however, as this interaction was not pronounced for any reproductive performance outcomes, it was excluded from the final models. Neonatal mortality was recorded as zero in most animals; therefore, GLMMs did not converge for this outcome, and only descriptive statistics (mean and range) were reported.

Behavioral and salivary cortisol data were analyzed using GLMMs that included housing treatment, experimental week, parity group, and their interactions (housing treatment × experimental week and housing treatment × parity group), with animal included as a random effect. A three-way interaction (housing treatment × experimental week × parity group) was not included due to the limited sample size. Additionally, the interaction between experimental week and parity group was excluded to avoid model overfitting, as it was not the primary focus of this study. Behavioral data were modeled using a binomial distribution with a logit link function, whereas salivary cortisol was modeled using a gamma distribution with a log link function.

All GLMMs and LMMs were implemented using the glmmTMB package. The significance of fixed effects was assessed using Type III Wald chi-squared tests via the *Anova* function in the *car* package. Post-hoc pairwise comparisons were conducted only for significant interactions using Tukey’s adjustment in the *emmeans* package. For the main effects of week and parity group, *p*-values from the Wald tests and estimated marginal means with 95% confidence intervals (CIs) were reported to describe overall trends. For presentation, estimated marginal means from GLMMs were back-transformed using the inverse link function. For behavioral indices, the response variable represented the proportion of sampling points in which the behavior was observed (out of 90 per day).

## RESULTS

No cases of lameness or abortion were observed in any sows throughout the experimental period.

The reproductive performance results are presented in Table 2. The number of stillbirths per litter was significantly lower in the PG group than in the CS group (*p* < 0.05; CS: 1.49 [0.86, 2.60] vs. PG: 0.63 [0.29, 1.36]). No cases of apparent infectious stillbirths were observed in this study. No significant effects of housing treatment or parity group were observed for total litter size, number of live piglets at the beginning of lactation, or average birth weight (Table 2). Neonatal mortality (defined as death before the onset of suckling due to crushing or weakness) occurred rarely; therefore, statistical analysis was not conducted, and the results are presented as mean (minimum–maximum) values (Table 2).

**Table 2.** Reproductive performance of sows assigned to continuous stall (CS) or periodic group housing systems.

|  | Treatment |  |  |  |  | Parity |  |  |  |  |  |  |
| --- | --- | --- | --- | --- | --- | --- | --- | --- | --- | --- | --- | --- |
|  | CS |  | PG |  | *P*-values | Young |  | Middle |  | Old |  | *P*-values |
| Total litter size | 16.07 | [14.09, 18.31] | 15.87 | [13.85, 18.19] | 0.90 | 15.93 | [13.82, 18.37] | 15.99 | [13.96, 18.32] | 15.98 | [12.83, 19.91] | 0.99 |
| Stillbirths | 1.49 | [0.86, 2.60] | 0.63 | [0.29, 1.36] | <0.05 | 1.10 | [0.58, 2.09] | 0.83 | [0.41, 1.67] | 1.00 | [0.37, 2.70] | 0.81 |
| Neonatal mortality <sup>1)</sup> | 0.20 | (0–3) | 0.47 | (0–4) | N.D. | 0.42 | (0–4) | 0.38 | (0–3) | 0.00 | (0–0) | N.D. |
| Live piglets at the beginning of lactation | 14.21 | [12.37, 16.33] | 14.73 | [12.79, 16.97] | 0.71 | 14.21 | [12.22, 16.52] | 14.56 | [12.62, 16.79] | 14.65 | [11.64, 18.44] | 0.96 |
| Average birth weight (kg) | 1.41 | [1.30, 1.51] | 1.34 | [1.23, 1.45] | 0.34 | 1.44 | [1.33, 1.55] | 1.45 | [1.34, 1.56] | 1.23 | [1.06, 1.41] | 0.08 |

Values in square brackets represent 95% CIs (lower and upper limits).
N.D., not determined; CS, continuous stall housing; PG, periodic group housing.
<sup>1)</sup> Due to zero inflation in the data, statistical analysis was not performed for neonatal mortality. Data are presented as means (minimum– maximum).

Behavioral indices and salivary cortisol concentrations are presented in Table 3, which reports estimated marginal means (95% CIs) for housing treatments, along with Wald test *p*-values for main effects and interactions. Estimated marginal means (95% CIs) for the main effects of experimental week and parity group are provided in Table 4.

**Table 3.** Estimated marginal means [95% CI] for the main effect of treatment and *p*-values for main effects and interactions on behavioral and physiological indices.

|  | Treatment |  |  |  | *P*-values |  |  |  |  |
| --- | --- | --- | --- | --- | --- | --- | --- | --- | --- |
|  | CS |  | PG |  | Treatment | Week | Parity | Trt × Week | Trt × Parity |
| Standing | 0.43 | [0.30, 0.58] | 0.17 | [0.10, 0.29] | ** | ** | 0.16 | 0.38 | * |
| Lying | 0.49 | [0.38, 0.61] | 0.76 | [0.65, 0.84] | ** | ** | * | 0.15 | ** |
| Sitting | 0.02 | [0.01, 0.06] | 0.01 | [0.00, 0.04] | 0.84 | ** | 0.70 | 0.52 | 0.42 |
| Drinking | 0.05 | [0.03, 0.08] | 0.01 | [0.01, 0.03] | † | † | † | 0.97 | 0.47 |
| Exploring | 0.28 | [0.18, 0.39] | 0.13 | [0.08, 0.22] | * | ** | 0.15 | † | 0.20 |
| Oral abnormal behavior | 0.39 | [0.26, 0.53] | 0.26 | [0.15, 0.40] | 0.39 | ** | 0.19 | ** | 0.14 |
| Cortisol (µg/mL) | 5.51 | [4.74, 6.40] | 4.75 | [4.02, 5.62] | 0.77 | ** | 0.25 | 0.78 | 0.48 |

Data are presented as estimated marginal means with 95% CIs. CS, continuous stall housing; PG, periodic group housing. Main effects and interactions were assessed using Wald chi-square tests based on generalized linear mixed models. \*\**p* < 0.01, \**p* < 0.05, †*p* < 0.1.

**Table 4.** Estimated marginal means [95% CI] for the main effects of week and parity on behavioral and physiological indices.

|  | Week |  |  |  |  |  | Parity |  |  |  |  |  |
| --- | --- | --- | --- | --- | --- | --- | --- | --- | --- | --- | --- | --- |
|  | Week1 |  | Week6 |  | Week10 |  | Young |  | Middle |  | Old |  |
| Standing | 0.33 | [0.24, 0.43] | 0.30 | [0.21, 0.40] | 0.24 | [0.16, 0.33] | 0.31 | [0.19, 0.46] | 0.30 | [0.19, 0.45] | 0.25 | [0.11, 0.47] |
| Lying | 0.61 | [0.53, 0.70] | 0.60 | [0.52, 0.69] | 0.69 | [0.61, 0.76] | 0.63 | [0.50, 0.73] | 0.57 | [0.45, 0.69] | 0.71 | [0.53, 0.84] |
| Sitting | 0.01 | [0.01, 0.03] | 0.02 | [0.01, 0.05] | 0.02 | [0.01, 0.04] | 0.02 | [0.01, 0.05] | 0.03 | [0.01, 0.07] | 0.01 | [0.00, 0.05] |
| Drinking | 0.02 | [0.01, 0.04] | 0.02 | [0.01, 0.04] | 0.03 | [0.02, 0.05] | 0.03 | [0.02, 0.05] | 0.05 | [0.03, 0.08] | 0.01 | [0.00, 0.03] |
| Exploring | 0.23 | [0.16, 0.30] | 0.19 | [0.13, 0.26] | 0.17 | [0.12, 0.24] | 0.25 | [0.16, 0.37] | 0.24 | [0.15, 0.36] | 0.12 | [0.05, 0.24] |
| Oral abnormal<br>behavior | 0.3 | [0.22, 0.40] | 0.37 | [0.27, 0.48] | 0.29 | [0.20, 0.38] | 0.39 | [0.26, 0.55] | 0.32 | [0.20, 0.47] | 0.25 | [0.11, 0.47] |
| Cortisol<br>(µg/mL) | 4.41 | [3.81, 5.11] | 4.87 | [4.22, 5.64] | 6.23 | [5.38, 7.21] | 5.49 | [4.68, 6.46] | 5.39 | [4.60, 6.32] | 4.52 | [3.52, 5.82] |

Data are presented as estimated marginal means with 95% CIs.

A significant interaction between housing treatment and parity group was observed for the proportions of standing and lying behaviors (Table 3). Post-hoc multiple comparisons showed no significant differences in standing behavior among groups (Fig. 2a). For lying behavior, a marked difference between housing treatments was observed only in the old parity group, where the proportion of lying behavior was higher in PG than in CS (Fig. 2b).

**Figure 2.**
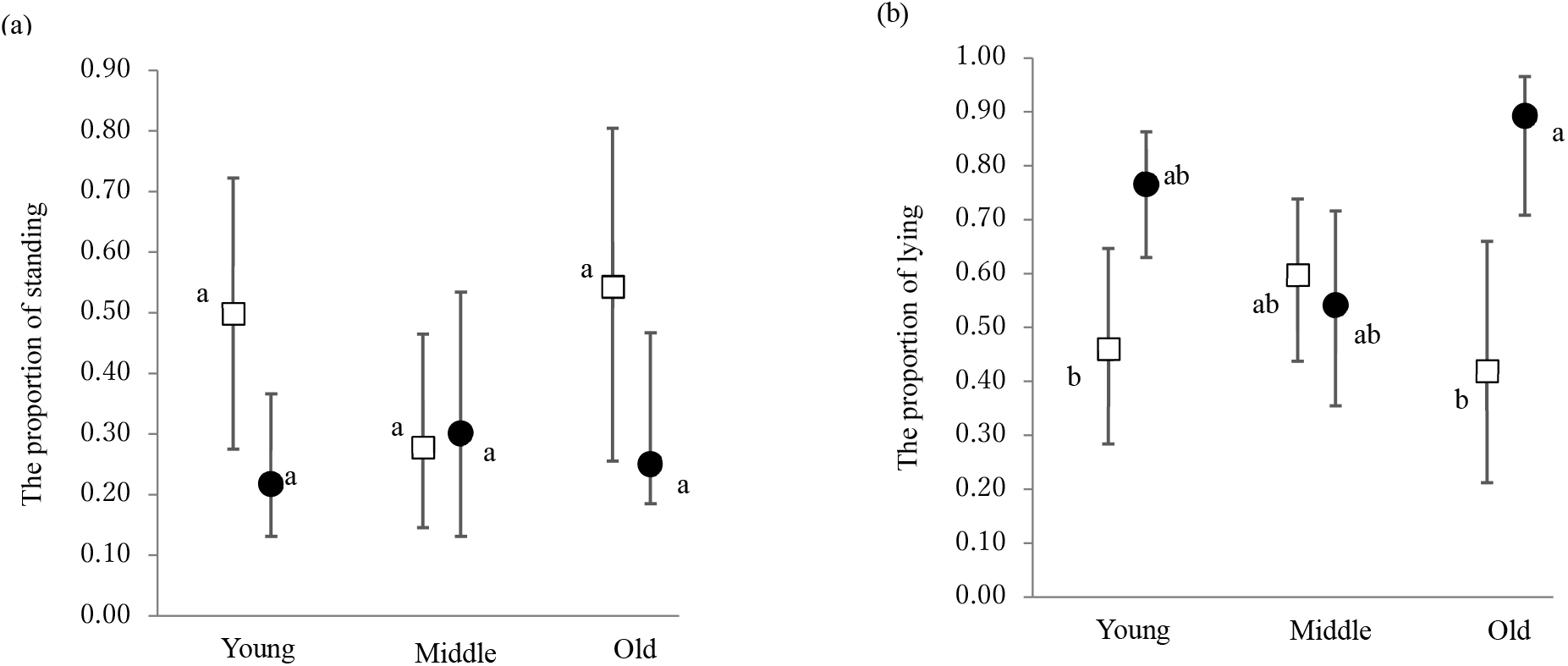
Effect of the interaction between housing treatment and parity group on the proportions of standing and lying. (a) Standing; (b) Lying. White squares indicate continuous stall housing (CS), and black circles indicate periodic group housing (PG). Error bars represent the lower and upper limits of the 95% CI. Different letters (a, b) indicate significant differences (*p* < 0.05).

The proportion of drinking behavior tended to be lower in PG than in CS (Table 3; CS: 0.05 [0.03, 0.08] vs. PG: 0.01 [0.01, 0.03]). The proportion of exploratory behavior was markedly lower in PG compared with CS (Table 3; CS: 0.28 [0.18, 0.39] vs. PG: 0.13 [0.08, 0.22]). No pronounced interactions involving housing treatment were detected for either behavior (Table 3).

A significant interaction between housing treatment and experimental week was observed for oral abnormal behavior (*p* < 0.01; Table 3). Post-hoc multiple comparisons revealed a significant increase from week 1 to week 6 in the PG group and a significant decrease from week 6 to week 10 in the CS group; however, no marked differences between housing treatments were detected within individual weeks (Fig. 3).

**Figure 3.**
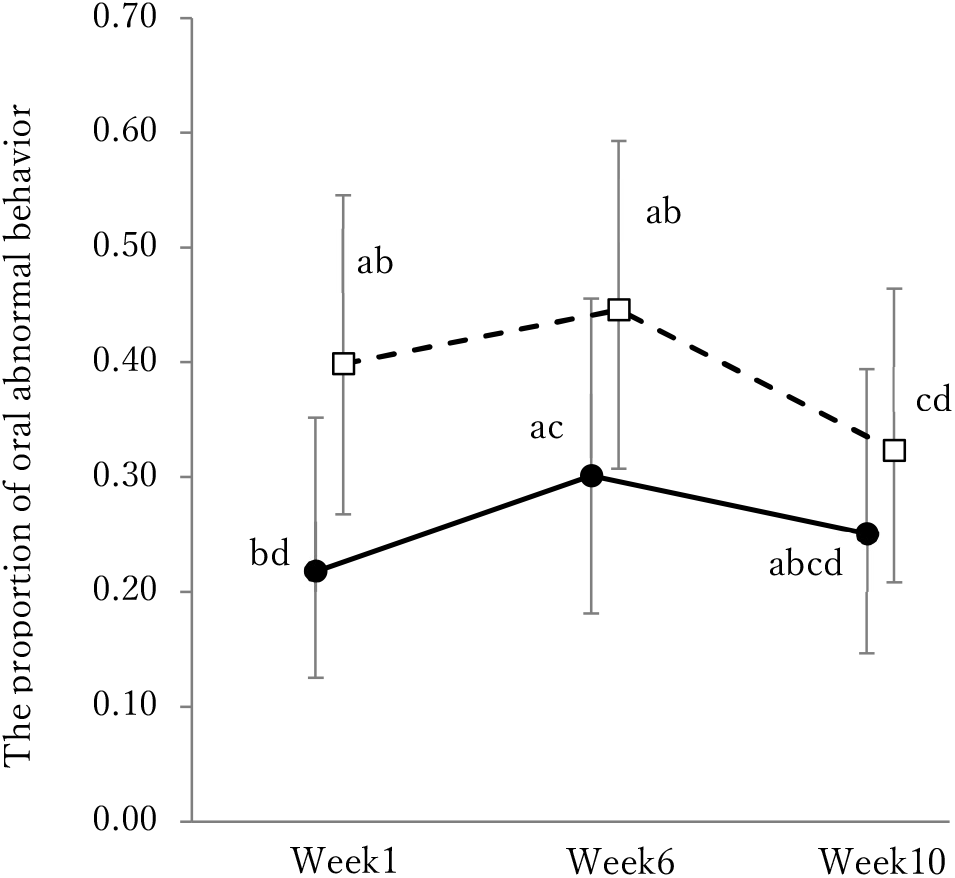
Effect of the interaction between housing treatment and experimental week on the proportion of oral abnormal behavior. White squares with a dotted line indicate continuous stall housing (CS), and black circles with a solid line indicate periodic group housing (PG). Error bars represent the lower and upper limits of the 95% CI. Different letters (a–d) indicate significant differences (*p* < 0.05).

Salivary cortisol concentrations did not differ between housing treatments, and no pronounced interactions involving housing treatment were detected (Table 3).

## DISCUSSION

### Effects on reproductive performance

In this study, PG was associated with fewer stillbirths per litter (CS: 1.49 vs. PG: 0.63), whereas total litter size was unaffected. Previous studies have reported that continuous group housing systems, such as electronic sow feeding systems or floor-feeding systems, reduce the number of stillborn piglets compared with individual stall housing, whereas total litter size remains unchanged [13–15]. Furthermore, Tokareva et al. (2022) reported that 10 min of weekly walking for individual stall-housed sows improved reproductive performance in older parity sows; however, the authors concluded that this limited intervention was insufficient to improve overall piglet survival [11]. Building on this finding, we evaluated a different intervention—weekly “group housing opportunities”— that allowed both free movement and social contact. A significant reduction in stillbirths was observed in the PG treatment, even after accounting for parity group. These results suggest that even relatively limited weekly access to group housing may reduce stillbirths in a manner broadly consistent with outcomes reported in continuous group housing systems.

No significant effects of housing treatment were observed for total litter size or average birth weight. This indicates that the PG treatment was not associated with measurable changes in fetal growth (average birth weight) or prenatal survival (total litter size), but may have reduced the risk of intrapartum mortality. The underlying mechanisms by which this weekly treatment significantly reduced stillbirths are likely related to the improved physical condition of the sow required for parturition and enhanced blood flow to the fetuses. Vanderhaeghe et al. (2013) reported that reduced physical activity due to prolonged confinement in stalls may lead to decreased uterine and overall muscle tone, which can impair the efficiency of expulsive efforts during parturition and is considered a major risk factor for increased stillbirths [16]. The primary cause of noninfectious stillbirths during parturition is fetal hypoxia associated with strong uterine contractions and prolonged farrowing [16,17]. Harris et al. (2013) reported that maternal exercise during gestation significantly increases umbilical blood flow [9]. Therefore, it is hypothesized that PG in this study may have improved sow muscular tone and uteroplacental and umbilical blood flow, thereby enhancing fetal tolerance to hypoxic stress during parturition. Further studies quantifying uterine physiological changes and farrowing duration are required.

### Effects on behavioral and physiological stress indicators

In this study, PG treatment significantly increased the proportion of time spent lying in stalls, especially in older parity sows. Chapinal et al. (2010) noted that restrictive environments such as stalls can induce frustration and stress in sows, thereby increasing overall arousal and activity levels [1]. Furthermore, stress associated with stall confinement has been reported to accumulate with increasing parity [18]. Accordingly, the lower proportion of lying observed in older parity sows under the CS treatment may reflect greater restlessness or reduced comfort under prolonged confinement. Conversely, the higher proportion of lying behavior among older parity sows in the PG treatment suggests that weekly opportunities for exercise and social contact may have reduced restlessness or improved comfort under stall housing.

The PG treatment significantly decreased exploratory behavior and tended to reduce drinking behavior across all parity groups. Pigs inherently possess strong and diverse motivations for exploration, including both foraging-related (extrinsic) and novelty-driven (intrinsic) behaviors [19,20]. In barren environments with limited feeding opportunities, these innate motivations are severely thwarted, leading to redirected exploratory behavior toward pen structures, such as floors or bars. Similarly, excessive drinking frequently observed in confined sows is not necessarily driven by physiological needs but by nonfunctional excessive drinking (i.e., stereotypic behavior such as drinker-pressing) associated with restricted feeding and frustration for oral manipulation [2,21]. The presence of such redirected or stereotypic behaviors is generally considered indicative of difficulty coping with the environment and reduced welfare [21]. Therefore, the higher proportions of exploratory and drinking behaviors observed in the CS treatment may reflect unfulfilled motivation for environmental exploration and oral manipulation. In contrast, weekly group housing sessions in the PG treatment may have partially satisfied motivations for exploration and oral manipulation, thereby reducing arousal upon return to stalls and decreasing redirected behaviors. Unlike postural behaviors (standing and lying), which showed a significant interaction between housing treatment and parity group, no such interaction was observed for exploratory or drinking behaviors. This indicates that the reductions in these behaviors associated with PG were consistent across parity groups and not restricted to older sows.

Oral abnormal behavior showed a significant interaction between housing treatment and experimental week. The significant increase from week 1 to week 6 in the PG group may be related to the relatively uniform environment provided during group housing sessions and to pigs’ tendency to respond more strongly to novelty than to familiarity [22]. Previous studies have shown that providing environmental enrichment, such as straw or wooden blocks, reduces sham-chewing in pregnant sows [23,24], suggesting that such materials can satisfy motivations for oral manipulation. Additionally, engagement with enrichment materials has been reported to persist longer when items are varied daily, consistent with pigs’ preference for novelty [25]. In this study, sawdust provided during group housing sessions may have served as an oral manipulation resource. Since the sows had no access to sawdust for at least 10 weeks before the experiment, the material was likely novel during week 1, which may explain the relatively low proportion of oral abnormal behavior at that time. However, repeated exposure may have led to habituation, reducing the novelty value of the material and resulting in increased abnormal behaviors observed in week 6.

In this study, behavioral observations were conducted on days when sows in the PG treatment had returned to their stalls. Nevertheless, an increase in lying behavior was observed in older parity sows, along with decreases in exploratory and drinking behaviors across all parity groups. In a previous study, Tokareva et al. (2022) reported that 10 min of individual walking exercise once per week did not improve lying behavior or reduce stereotypic behaviors in stall-housed sows [26]. In contrast, the PG treatment in this study provided a prolonged opportunity for exercise of approximately 24 h, in addition to allowing social contact with others. These findings suggest that longer duration outside stalls and the availability of social contact may have helped to satisfy behavioral motivations for movement, exploration, and oral manipulation, thereby contributing, at least in part, to sustained comfort after the animals returned to stall housing.

No differences in salivary cortisol concentrations were detected between treatments. This may be related to the diurnal rhythm of cortisol and the limitations of this measure in assessing chronic stress [1]. It is well established that cortisol concentration shows a clear diurnal rhythm with high levels in the early morning in pregnant sows [27]. Furthermore, chronic stress or barren housing conditions sometimes attenuate this circadian rhythm rather than simply increasing the overall baseline [28]. In this study, salivary samples were collected once per sampling day between 08:30 and 09:00 h, a time period associated with relatively high baseline cortisol concentrations [27]. Consistent with this, Tatemoto et al. (2019) reported no differences in morning salivary cortisol levels between barren- and enriched-housed pregnant sows, whereas afternoon concentrations were significantly lower in the enriched group [24]. Hence, to more accurately evaluate the effects of PG on cortisol dynamics, additional sampling at times of lower baseline cortisol levels, such as late afternoon, may be required.

Although the present findings suggest potential benefits of PG, several limitations should be considered. The number of animals in each treatment by parity group was limited, particularly in the old parity group; therefore, the parity-specific increase in lying behavior observed in older sows should be interpreted cautiously. Moreover, the PG treatment consisted of several inseparable components, including exercise, social contact, and access to sawdust in the group housing area. The present study also did not examine how variations in group size, the frequency of group housing opportunities, or the characteristics of the group housing area, such as space allowance and the type of enrichment materials, might influence the effects of PG. The findings should therefore be interpreted as reflecting the combined effects of the PG treatment under the specific conditions of the present study. Future studies should include larger numbers of sows, especially those in the old parity group, and experimental designs that can separate the effects of the treatment components and housing conditions described above. Additionally, behavioral observations during group housing sessions, direct measurements of farrowing processes related to stillbirth, and repeated or continuous physiological measurements would help clarify the mechanisms underlying the effects of PG.

In conclusion, under the conditions of this study, PG showed several potential benefits. First, the number of stillbirths per litter was significantly reduced, whereas the total litter size was unaffected. Second, the proportion of lying behavior significantly increased in older parity sows, which may indicate improved comfort under conditions of restricted movement. Third, the proportion of exploratory behavior significantly decreased, and drinking behavior showed a decreasing trend, suggesting partial satisfaction of motivations for environmental exploration and oral manipulation. Although further refinement of the system, such as rotating enrichment materials, may be required to reduce specific stereotypic behaviors more effectively, these findings indicate that the PG system may contribute to improved reproductive performance and specific measurable aspects of sow welfare.

## CONFLICT OF INTEREST

The authors declare that there are no conflicts of interest with any financial organization regarding the study reported in this manuscript.

## AUTHORS’ CONTRIBUTIONS

Tomoya Shimasaki, Ken-ichi Yayou, and Ken-ichi Takeda conceptualized and designed the experiment. Tomoya Shimasaki, Ken-ichi Yayou, Tomoki Kojima, and Chen-Yu Huang conducted the animal experiments. Tomoya Shimasaki, Ken-ichi Yayou, Tomoki Kojima, Mitsuyoshi Ishida, and Ken-ichi Takeda analyzed the data and interpreted the results. All authors contributed to the revision of the manuscript and approved the final version for publication.

## FUNDINGS

This study was supported by the MAFF Commissioned project study on “Development of comfortable management systems for poultry and swine” (Grant No. JPJ011279) and the Ito Foundation (Grant No. 2024-79).

## ACKNOWLEDGMENTS

The authors gratefully acknowledge the members of the Medium Livestock Technical Team, Ikenodai Operation Unit 2, Technical Support Office, NIGLS, NARO, for their assistance with this study. The authors also thank Ms. Miyoko Arai for her support in preparing and cleaning the experimental equipment.

## DATA AVAILABILITY

The datasets generated and/or analyzed during this study are available from the corresponding author upon reasonable request.

## DECLARATION OF GENERATIVE AI

During the preparation of this manuscript, the authors used Microsoft Copilot to assist with improving readability and grammatical accuracy. After the use of this tool, the authors have reviewed and edited the content as necessary and accepted full responsibility for the content of the manuscript.

## Notes

### Competing Interest Statement

The authors have declared no competing interest.

